# Simultaneous estimation of transcript abundances and transcript specific fragment distributions of RNA-Seq data with the Mix^2^ model

**DOI:** 10.1101/005918

**Authors:** Andreas Tuerk, Gregor Wiktorin

**Affiliations:** Lexogen GmbH, Campus Vienna Biocenter 5, 1030 Vienna, Austria

## Abstract

Quantification of RNA transcripts with RNA-Seq is inaccurate due to positional fragmentation bias, which is not represented appropriately by current statistical models of RNA-Seq data. Another, less investigated, source of error is the inaccuracy of transcript start and end annotations.

This article introduces the Mix^2^ (rd. “mixquare”) model, which uses a mixture of probability distributions to model the transcript specific positional fragment bias. The parameters of the Mix^2^ model can be efficiently trained with the EM algorithm and are tied between similar transcripts. Transcript specific shift and scale parameters allow the Mix^2^ model to automatically correct inaccurate transcript start and end annotations. Experiments are conducted on synthetic data covering 7 genes of different complexity, 4 types of fragment bias and correct as well as incorrect transcript start and end annotations. Abundance estimates obtained by Cufflinks 2.2.0, PennSeq and the Mix^2^ model show superior performance of the Mix^2^ model in the vast majority of test conditions.

The Mix^2^ software is available at http://www.lexogen.com/fileadmin/uploads/bioinfo/mix2model.tgz, subject to the enclosed license.

Additional experimental data are available in the supplement.

## 1 Introduction

In recent years RNA-Seq has established itself as a popular alternative to microarrays for the quantification of RNA transcripts. In contrast to microarrays, which measure the quantity of an RNA transcript by hybridization to a transcript specific oligonucleotide, RNA-Seq generates cDNA fragments for sections of the RNA transcript, which are sequenced by a next generation (NGS) sequencer. One advantage of RNA-Seq over microarrays is that it does not require prior knowledge of the nucleotide sequence of the RNA transcript, which is needed to fashion a transcript specific hybridization probe, and that it can therefore detect and quantify novel RNA transcripts. In addition, quantification by RNA-Seq covers a wider dynamic range since microarrays suffer from signal saturation resulting in the truncation of abundance estimates for highly abundant transcripts [4].

Despite these advantages, obtaining accurate quantification measurements from RNA-Seq has proven difficult. This is in part due to the technical variability of RNA-Seq protocols but also a result of the oversimplified representation of RNA-Seq data by the statistical models used to obtain quantification measurements. In particular, quantification methods, such as Cufflinks [15], assume that the cDNA fragments generated by RNA-Seq are distributed evenly across the transcripts. This, however, contradicts the majority of RNA-Seq data which exhibit biased rather than uniform fragment distributions and the above assumption therefore leads to inaccurate transcript quantification. In order to improve transcript quantification it is important to avoid unrealistic assumptions regarding the nature of fragment bias and to modify the statistical models in a way which enables them to automatically adapt to the bias present in RNA-Seq data.

One type of bias affecting transcript quantification based on RNA-Seq is the result of a preference of the fragmentation, i.e. the process that generates cDNA fragments from RNA transcripts, to produce fragments at certain positions within the transcript, e.g. at the start and/or at the end of the transcript. Hence, this type of bias is referred to as positional bias [14]. Positional bias can also be caused by a bias in the RNA itself, for instance, due to RNA degradation which results in a shortening of the RNA. Another kind of bias in RNA-Seq is introduced during ligation, amplification and NGS sequencing [3]. This bias is correlated to the RNA sequence of a transcript and is therefore termed sequence specific bias [14]. The present article focuses on the first type of bias, i.e. the positional bias, and develops a model, the Mix^2^ model, which is shown to perform significantly better than both Cufflinks [15] and PennSeq [5]. Furthermore, as shown in Section 2 of the supplement, the Mix^2^ model can easily be combined with other bias models, such as the fragment specific model in [14] to obtain a complete model for all observed biases.

Another, less investigated, source of error in the quantification of RNA transcripts with RNA-Seq is the inaccuracy of the transcript start and end annotations. This inaccuracy might, for instance, stem from the start and end site variation of RNA transcripts [12]. In addition to modelling positional bias, the Mix^2^ model can also correct inaccurate transcript start and end annotations automatically and in this situation therefore performs substantially better than the compared methods.

The inclusion of bias models into the statistical models of RNA-Seq data has been investigated before. [9] proposes a model to account for the variability in read counts depending on the sequence surrounding the start of a fragment. The intention is similar to that of the fragment specific bias model in [14], which also investigates the sequences surrounding the start of a fragment. Similar to [15], the generative models in [7] and [6] describe the probability of observing a fragment for a given transcript. In addition to [15], however, [7] and [6] introduce additional hidden variables and use a bias model, which is derived from the global bias observed in the complete RNA-Seq data set. The method described in [17], on the other hand, is a model for gene read counts, which models bias by exon specific weights, which are estimated both for the complete data set and for individual genes. [16] focuses on RNA-Seq data with 5’ bias which is the result of RNA degradation and uses an exponential model for the fragment distributions. The model proposed in [10] is, again, a model for the read counts of a gene. Here the read counts are modelled by a quasi-multinomial distribution with a parameter that can be adapted to account for over and under dispersion. [14] proposes a model both for sequence specific and positional bias. For the positional bias a non-parametric model is used, which can theoretically be trained with the EM algorithm. However, due to the large number of variables, this is only feasible for a drastically simplified version of the model. One of the simplifications is that the positional bias model in [14] is only trained on genes containing a single transcript. The model developed in [5] is again non-parametric and the large number of variables makes training the full model computationally prohibitive. As in [14], the model in [5] needs to be simplified considerably in order to obtain a feasible adaptation procedure.

Similar to some of the other methods mentioned above, the Mix^2^ model is a generative model for the probability of observing a fragment for a given transcript. In contrast to the positional bias model in [14] and PennSeq [5], however, the Mix^2^ model is parametric, which considerably simplifies adaptation of the Mix^2^ parameters with the EM algorithm. On the other hand, due to the representation of the transcript specific fragment bias by mixtures of probability distributions, the Mix^2^ model is flexible enough to allow for positional biases of arbitrary complexity.

## 2 Methods

### 2.1 Background

An essential step in next generation sequencing (NGS) of transcriptomes is the library preparation. This process takes an RNA sample and produces a library of short cDNA fragments, each corresponding to a section of an RNA transcript. The cDNA fragments are sequenced by an NGS sequencer resulting in single or paired end reads which are mapped to a reference genome. In the case of paired end reads, the start and end position of the original fragment can thus be located in genome coordinates. In the following it will therefore be assumed that for a fragment *r* the start and end of the fragment can be inferred from the data.

The distribution *p*(*r*) of the fragments *r* in a genomic locus is a superposition of the fragment distributions *p*(*r*|*t* = *i*) for the *N* transcripts in the locus, i.e.

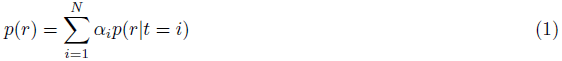

where *α_i_* is the abundance of transcript *t* = *i*, i.e. the probability that transcript *t* = *i* generates any fragment, and *p*(*r*|*t* = *i*) is the probability that transcript *t* = *i* generates fragment *r*. An estimate for the concentration of transcript *t* = *i* is obtained by normalizing the abundance *α_i_*, yielding the RPKM [11] or FPKM values [15].

Given a set of fragments *R* in a genomic locus and the fragment distributions *p*(*r|t* = *i*), the transcript abundances can be estimated by maximizing the likelihood of *R* with the expectation maximization (EM) algorithm [1]. For the *n* + 1th iteration of the EM algorithm this yields the following formula

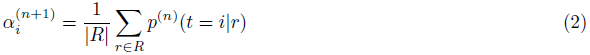

where *p*^(^*^n^*^)^(*t* = *i*|*r*) is the posterior probability of transcript *t* = *i* given fragment *r* after *n* iterations of the EM algorithm and 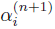 is the estimate for *α_i_* after *n* + 1 iterations.

The fragment distributions *p*(*r|t* = *i*) are not known in an RNA-Seq experiment and have therefore either to be estimated from the data or approximated by an appropriate statistical model. In Cufflinks [15], for instance, the *p*(*r*|*t* = *i*) are factorized as follows

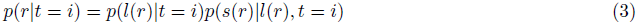

where *s*(*r*) and *l*(*r*) are the start and length of fragment *r* in the coordinates of transcript *t* = *i*. The fragment length distribution *p*(*l*(*r*)|*t* = *i*) is derived from the cumulative statistic of fragment lengths by truncation and renormalization. The fragment start distribution *p*(*s*(*r*)|*l*(*r*)*, t* = *i*), on the other hand, is assumed to be uniform over the possible fragment starts *s*(*r*) for transcript *t* = *i* and fragment length *l*(*r*). For a cumulative fragment length distribution given by a Gaussian with mean 200 bp and standard deviation 80 bp, the fragment start distributions *p*(*s*(*r*)|*t* = *i*) of the Cufflinks model, which are derived from (3) by summing out *l*(*r*), are visualized in Figure 3(a) for transcripts with lengths between 400 bp and 3000 bp. Figure 3(a) shows that the fragment start distributions of the Cufflinks model are unbiased for transcripts above a certain length which is an unrealistic assumption for most RNA-Seq data leading to inaccurate abundance estimates.

### 2.2 The Mix^2^ model

In contrast to the static model of Cufflinks, the model developed in this article estimates the fragment distributions *p*(*r*|*t* = *i*) from the RNA-Seq data. This is achieved by using a mixture model for *p*(*r*|*t* = *i*), i.e.

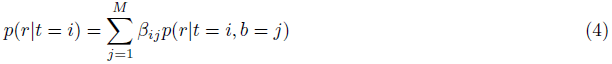

where *M* is the number of mixture components and the hidden variable *b* = *j* represents a “building block” of the fragment distribution *p*(*r*|*t* = *i*). In addition, for each *i*, the *β_ij_* sum to one. Since *p*(*r*) is itself a mixture of the *p*(*r|t* = *i*) with weights *α_i_*, this implies that *p*(*r*) is a mixture of mixtures and hence the model developed in the following will be called the Mix^2^ (pr. “mixquare”) model. In addition to the *α_i_* and *β_ij_*, the Mix^2^ model can depend, as in Section 2.2.2, on further internal parameters of the *p*(*r|t* = *i, b* = *j*).

If the distribution of the fragment length *l*(*r*) depends only on *s*(*r*) and is independent of *b* = *j* then (4) reduces to

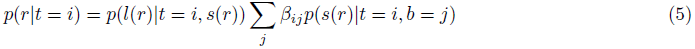

In this case, the *β_ij_* parameters model the positional variation of the fragment start *s*(*r*). An example for the Mix^2^ model in (5) representing data with a 5’ bias is given in Figures 1(a) and 1(b). Figure 1(a) shows a set of 8 *β_ij_*, whereas the dashed curves in Figure 1(b) show the mixture components *p*(*s*(*r*)|*t* = *i, b* = *j*) multiplied by the *β_ij_* for a transcript of 2000 bps length. The *p*(*s*(*r*)|*t* = *i, b* = *j*), in this case, are Gaussians with a constant standard deviation of 2000*/*18 and equidistant means *µ_j_* = *j* · 2000*/*9 normalized to the 2000 possible values of *s*(*r*). The solid curve in Figure 1(b) is the sum of the dashed curves and thus equals *p*(*r|t* = *i*). Hence, the *p*(*r|t* = *i*) in Figure 1(b) models a fragment distribution with a 5’ bias.

**Figure 1:**
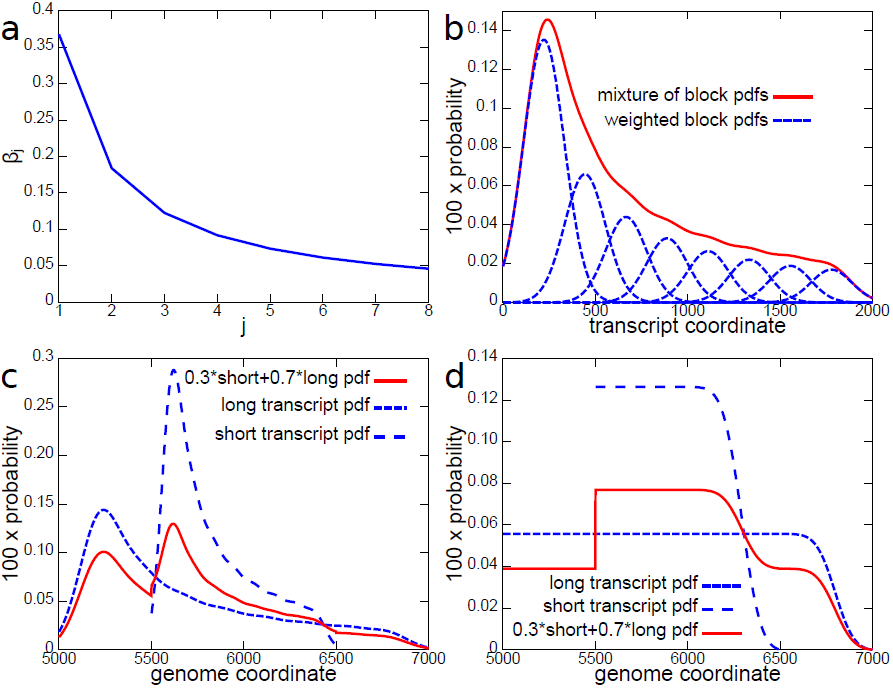
Fragment start distributions modelled by the Mix^2^ and the Cufflinks model. With the *β_j_* in the left upper graph the Mix^2^ model represents the fragment start distribution in the right upper graph of the figure. For the two transcripts in the bottom left graph, which share the same set of *β_j_* and for which *α*_1_ = 0.7 and *α*_2_ = 0.3 the Mix^2^ model represents the fragment start distribution shown by the red line. In comparison, the Cufflinks model can only model loci in which the fragment start distribution is similar to the distribution in the bottom right graph.

The parameters of the Mix^2^ model are estimated by maximizing the likelihood of the set of fragments *R* in a genomic locus, with the goal of obtaining a simultaneous estimate for the abundances *α_i_* and for the fragment distributions *p*(*r|t* = *i*). Optimizing the parameters of the Mix^2^ model without any further restrictions leads to a good fit of the set of fragments *R* by the distribution *p*(*r*), but not necessarily to a correct estimate of the abundances *α_i_* or the fragment distributions *p*(*r*|*t* = *i*). In order to achieve this task, similarities between the *p*(*r*|*t* = *i*) for different transcripts *t* = *i* have to be included into the Mix^2^ model. This can be done by parameter tying. In the case of the Mix^2^ model in (5), for instance, the *β_ij_* can be tied for transcripts whose fragment starts *s*(*r*) have a similar positional bias. The topic of parameter tying, which is essential for the Mix^2^ model, will be discussed in the following three sections.

#### 2.2.1 Tying the mixture weights of the Mix^2^ model

A straight forward type of parameter tying in the Mix^2^ model is the tying of the *β_ij_*between different transcripts *t* = *i*, i.e. *β_ij_* = *β_j_* and hence

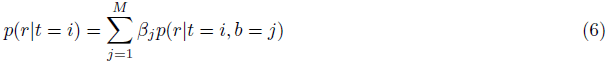

This together with (1), after summing out the *t* = *i*, implies the following

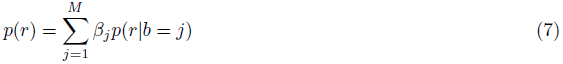

Comparing (7) and (1) shows that, given the abundances *α_i_*, the *β_j_* can be estimated, similarly to (2), with the EM algorithm as follows

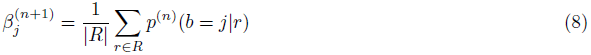

Unlike the individual optimization of *α_i_* and *β_j_*, the simultaneous optimization of *α_i_* and *β_j_* is not a linear problem and local maxima therefore exist. In practice, however, the difference between the *α_i_* and *β_j_* at different local maxima was found to be negligible and in the experiments in section 3 it was therefore sufficient to run the EM algorithm with a single parameter initialization, in order to obtain reliable estimates of *α_i_* and *β_j_* from (2) and (8).

Figure 1 shows an example for a Mix^2^ model with tied *β_ij_* in a genomic locus containing two transcripts. The first transcript has abundance *α*_1_ = 0.7 is 2000 bps long and starts at the 5000’th bp of a contig in the genome. The second transcript has abundance *α*_2_ = 0.3 is 1000 bps long and starts at the 5500’th bp of the same contig. Neither transcript contains any introns and both share the same set of *β_j_* as in Figure 1(a). The distribution *p*(*s*(*r*)|*t* = *i*) in transcript coordinates for the long transcript is the distribution in Figure 1(b), whereas the distribution of the short transcript is derived by multiplying the *β_j_* in Figure 1(a) with Gaussians with a constant standard deviation of 2000*/*36 and means placed equidistantly across the coordinates of the short transcript, i.e. *µ_j_* = *j* 2000*/*18. The distributions *p*(*s*(*r*)|*t* = *i*) in genomic coordinates are visualized by the dashed lines in Figure 1(c) whereas *p*(*s*(*r*)) is given by the solid line. Thus, the Mix^2^ model in this example models a fragment start distribution *p*(*s*(*r*)) in the genomic locus where both transcripts have a similar 5’ bias. The static Cufflinks model, on the other hand, can only model the fragment distribution *p*(*s*(*r*)) in Figure 1(d), which is unbiased.

#### 2.2.2 Tying the parameters of the continuous Mix^2^ model

The distributions *p*(*r|t* = *i, b* = *j*) in Figure 1 are normalized so that their sum over the 2000 possible fragment starts equals one. This represents a finite probability space which incorporates the information that fragments can only start at a finite set of points and can only have a finite set of lengths. While this property is clearly shared by all fragments generated from a transcript, enforcing this requirement is inconvenient from a computational point of view. Instead, in the following, the probability distribution *p*(*r|t* = *i, b* = *j*) will be assumed to be a Gaussian on the continuous one or two dimensional probability space, i.e.

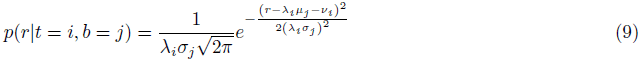

Here, *µ_j_* and *σ_j_* are parameters that are specific to block *b* = *j* and are shared between different transcripts, while *ν_i_* and *λ_i_*are parameters specific to transcript *t* = *i* and are shared for different blocks. The parameters *ν_i_*and *λ_i_* shift and scale the distribution *p*(*r*|*t* = *i*), respectively. Thus *ν_i_* and *λ_i_* can be used to position the fragment distribution *p*(*r*|*t* = *i*) at the correct start and end of the transcript *t* = *i*. In comparison to Figure 1 the *µ_j_* in (9) do not have to be placed at equidistant positions along the transcript coordinates and the *σ_j_* do not have to be constant. This additional degree of freedom allows for a more accurate modelling of the fragment distributions *p*(*r*|*t* = *i*) and a potentially smaller number of blocks in the mixture (4).

Using the continuous Gaussian (9) has the advantage that the internal parameters of the *p*(*r|t* = *i, b* = *j*), i.e. the *µ_j_, σ_j_, ν_i_, λ_i_*, can be estimated within the Mix^2^ model by the EM algorithm. For *ν_i_*, for instance, this yields the following update formula.

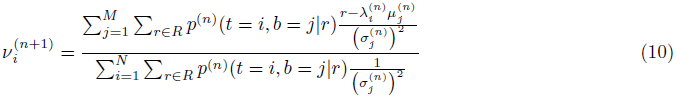

The derivation and update formulae for the remaining parameters *µ_j_, σ_j_* and *λ_i_* can be found in Section 1.1 in the supplement.

In order to correct transcript start and end annotations with the parameters *ν_i_* and *λ_i_* of the Mix^2^ model the original transcript annotation has to be extended. This process is visualized in Figure 2, which shows the initial, incorrect annotation as the thickest bar. The correct annotation with a transcript start upstream of the incorrect start and a transcript end downstream of the incorrect end is visualized by the bar of medium thickness. The extended annotation is given by the thin bar. It is necessary to extend the incorrect annotation, because otherwise, fragments are considered not to be compatible with the transcript due to the incorrect and shorter annotation. On the other hand, it is necessary to position *p*(*r*|*t* = *i*) within the extended annotation in such a way that *p*(*r*|*t* = *i*) = 0 for fragments *r* which are compatible with the extended but incompatible with the correct annotation. In Figure 2, for instance, the task of positioning *p*(*r*|*t* = *i*) correctly is achieved by setting *ν_i_* = 300 and *λ_i_* = 1.3 which places *p*(*r|t* = *i*) 300 bps from the start of the extended annotation and extends the length of the incorrect annotation to the correct transcript length.

**Figure 2:**
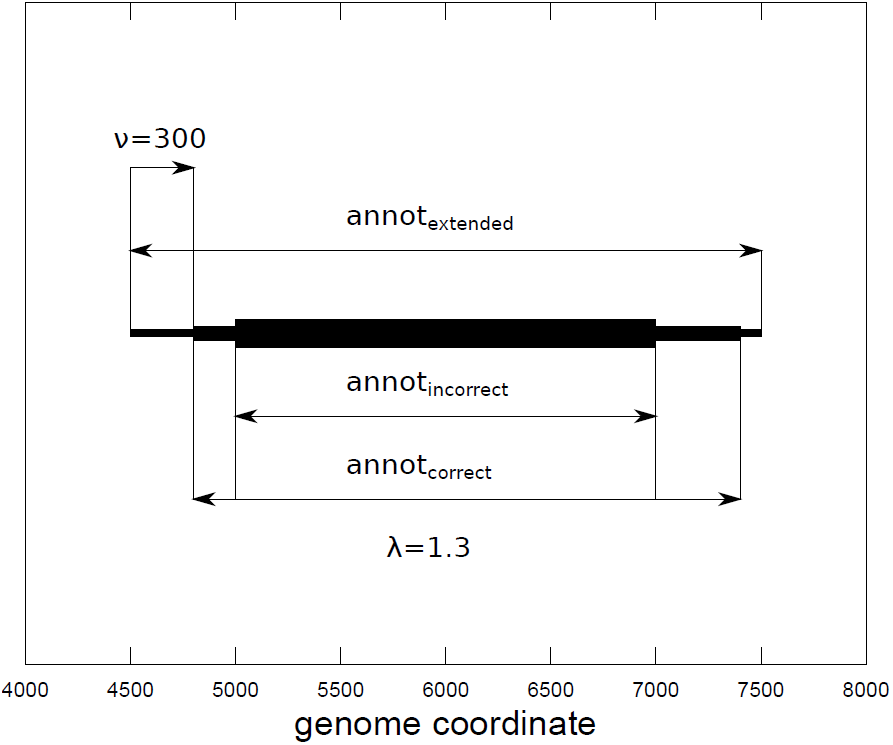
The continuous Mix^2^ model can correct incorrect transcript start and end annotations. In order to allow for the correction of an annotation whose end points lie outside the incorrect annotation, the incorrect annotation is extended. The parameters *ν* and *λ* are then used to position the transcript start and length correctly inside the extended annotation.

In comparison to the optimization of the weight parameters *β_j_* in section 2.2.1, the optimization of the parameters of the continuous Mix^2^ model is more strongly affected by the presence of local maxima in the likelihood function. For this reason, it is necessary to try out various initializations of the Mix^2^ model parameters in the EM algorithm to find good solutions. In the experiments in section 3.3, 10 random initializations were found to be sufficient to obtain good results.

#### 2.2.3 Tying the parameters of the Mix^2^ model within groups

Tying parameters between the fragment distributions of different transcripts results in similar *p*(*r*|*t* = *i*). If the *β_ij_* are tied as in Figure 1, for instance, each *p*(*s*(*r*)|*t* = *i*) can be derived from any other *p*(*s*(*r*)|*t* ≠ *i*) by scaling *p*(*s*(*r*)|*t* ≠ *i*) to the length of *t* = *i*. Such similarities might, however, not hold for all the transcripts within a genomic locus. Consider, for example, the fragment start distributions *p*(*s*(*r*)|*t* = *i*) of the Cufflinks model. These are given in Figure 3(a) for transcripts of 400, 700, 1000, 2000 and 3000 bp length. The x-axis in in this figure shows the relative position of the fragment start within the transcript in percent and the distributions are scaled such that the integral under each curve in Figure 3(a) equals one. The distributions *p*(*s*(*r*)|*t* = *i*) in Figure 3(a) for transcripts between 2000 and 3000 bps length are roughly similar and can therefore use a single set of *β_ij_*. The same holds true for transcripts between 700 and 1000 bps length. The distribution of transcripts of 400 bp length, on the other hand, is rather different from any other distribution and should therefore use its own set of *β_ij_*. If transcripts of such different lengths as in Figure 3(a) are present in a single genomic locus, this suggests that the transcripts should be divided into 3 or more groups of similar transcripts such that only the transcripts within each group share the same set of *β_ij_*. This leads to the scenario, where each transcript *t* = *i* has an associated group *g* = *k* and the distributions *p*(*r*|*t* = *i*) of transcripts within this group share a certain set of parameters, for instance the *β_ij_*. For the *j*-th *β* in group *g* = *k* the EM update formula is then given as follows.

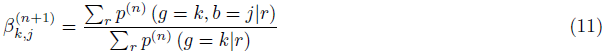

**Figure 3:**
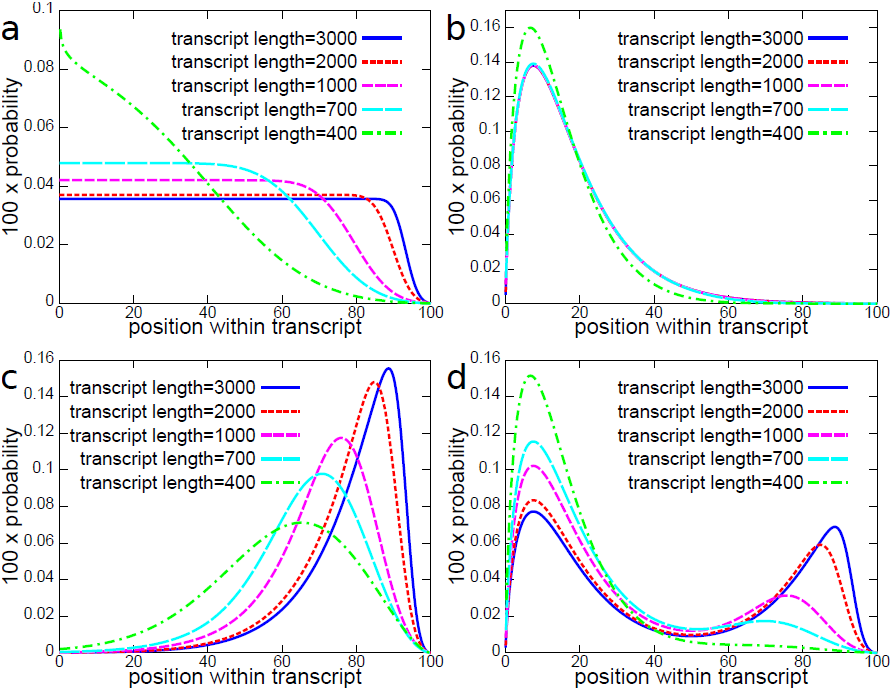
Fragment start distributions, from which data were sampled, for transcripts of 400, 700, 1000, 2000 and 3000 bps length. The x-axis is the position within the transcript in percent. The distributions are derived from an initial distribution which is scaled to the transcript length. Subsequently, this distribution is multiplied with the fragment start distribution *p*(*s*(*r*)|*t* = *i*) of Cufflinks and renormalized. Figure (a) shows the resulting distributions for a uniform initial distribution, which therefore corresponds to the fragment start distributions of Cufflinks. Figure (b), (c) and (d) show the distributions derived from an initial distribution with 5’ bias (b), 3’ bias (c) and 5’+3’ bias (d).

The derivation of (11) can be found in Section 1.2 in the supplement.

Placing each transcript in a genomic locus into its own group removes all parameter tying and leads, as mentioned before, to inaccurate estimates for fragment distributions and transcript abundances. When placing transcripts into different groups it is therefore important to strike the right balance between separation of dissimilar fragment distributions and retaining the tying of a sufficient number of parameters to ensure the stability of the Mix^2^ model.

## 3 Results

The experiments in this section compare the Mix^2^ model to Cufflinks 2.2.0 (http://cufflinks.cbcb.umd.edu/index.html) and the PennSeq abundance estimation in the version of 03/25/2014 (http://sourceforge.net/projects/pennseq). Cufflinks and PennSeq were chosen because Cufflinks [15] is one of the most widely used methods for the estimation of transcript abundances, while PennSeq [5] has recently been shown to outperform a large number of publicly available abundance estimation methods, including the bias models proposed in [14], [10] and [16].

The test data in the experiments cover 4 types of fragmentation bias and extensive sets of abundances on 7 test genes. Since the strand of the genes is irrelevant in the experiments, all genes were considered to lie on the plus strand. Similar to a recent review of computational RNA-Seq methods in [13], the test data in the experiments are exclusively synthetic and the correct abundances are therefore known. In order to gain better control over the fragment bias of the test data, rather than using the Fluxsimulator [2], the fragments in the experiments are directly sampled from the bias models.

The parameters of the Mix^2^ model are trained with the EM algorithm, which was terminated if the change in likelihood of the sampled fragments or the change of the parameters between successive iterations fell below a certain threshold.

The difference between true and estimated abundances is measured with the *L*_1_ distance, which is the sum of the absolute differences between true and estimated abundances and can therefore be interpreted as the accumulated error over the complete gene. The *L*_1_ distance of two probability distributions lies between 0 and 2.

### 3.1 Test data

Experiments were performed on a set of 7 genes from the human reference GRCh37 v73 of Ensembl. The genes were selected to cover abundance estimation scenarios of different complexity and contain between 4 and 15 transcripts as well as the main variants of differential splicing. The names and the properties of these genes are summarized in Table 1. With the exception of mutually exclusive exons all types of differential splicing are present. For each of the 7 genes, 200 sets of abundances were sampled uniformly, according to the Dirichlet distribution, from the simplex of probability distributions on the gene transcripts. Subsequently, for each of the 200 sets of abundances 10,000 fragments were sampled from the superposition in (1), where the *p*(*r|t* = *i*) belong to one of 4 different transcript length dependent bias models. Hence, for each abundance estimation method 800 experiments were performed per gene and 1400 experiments per bias model.

**Table 1:** The 7 test genes from the human reference GRCh37 v73 of Ensembl and their variants of differential splicing. The first three columns contain the number of transcripts (TR) within the gene and the number of base pairs of the longest (LT) and shortest transcript (ST). The remaining columns indicate which types of differential splicing are present, i.e. exon skipping (ES), alternative 5’ donor sites (A5DS), alternative 3’ acceptor sites (A3AS) intron retention (IR).

| gene | TR | LT | ST | ES | A5DS | A3AS | IR |
| --- | --- | --- | --- | --- | --- | --- | --- |
| DAPK3 | 7 | 2257 | 599 |  |  | × | × |
| HAUS5 | 10 | 4462 | 526 | × | × | × | × |
| KLK5 | 5 | 1563 | 672 | × |  | × | × |
| LDHD | 4 | 2067 | 701 |  | × | × | × |
| LGALS17A | 6 | 2584 | 406 | × |  | × | × |
| TESK2 | 7 | 3074 | 428 | × |  |  |  |
| USF2 | 15 | 2742 | 446 | × |  | × | × |

Figure 3 shows the four different types of transcript length dependent fragment start distributions *p*(*s*(*r*)|*t* = *i*) which were studied in the experiments. The first set of distributions in Figure 3(a) shows the fragment start distributions for the Cufflinks model for a fragment length distribution with mean 200 bps and standard deviation 80 bps. These distributions will therefore be referred to as Cufflinks bias in the following. Cufflinks 2.2.0 has to outperform PennSeq and the Mix^2^ model on fragments with Cufflinks bias and verification of this statement therefore serves as a control experiment which guarantees the integrity of the generated data.

The fragment start distributions in Figures 3(b), (c) and (d) were derived from the distributions in Figure 3(a) by multiplication with a biased distribution and subsequent renormalization. The fragment start distributions in Figure 3 therefore reflect the fact that for short transcripts fragments are predominantly located at the start of the transcript and the fragment start distribution therefore exhibits a 5’ bias. The bias models which were multiplied with the Cufflinks distributions in Figure 3(a) are distributions with a 5’ bias (b), 3’ bias (c) and a 5’+3’ bias (d). The biased distributions were derived from a Chi^2^ model by truncating the Chi^2^ distribution at the end of the transcript. For the 3’ bias in Figure 3(c) the truncated Chi^2^ distribution was mirrored and for the 5’+3’ bias in Figure 3(d) the 5’ and 3’ bias distributions were added each with a weight of 0.5. Thus, similar to the Cufflinks bias, for long transcripts the 5’+3’ bias distributions are almost symmetric. For this reason, Cufflinks bias and 5’+3’ bias are more similar than any other pair of biases and Cufflinks 2.2.0 can therefore be expected to perform well on data with 5’+3’ bias.

For the distributions with 5’ bias in Figure 3(b) the effect of the transcript length on the shape of the distributions is weak. For transcripts with a length between 700 and 3000 bps the shapes are almost identical. For the other biases, however, this effect is comparatively strong and using multiple groups in the Mix^2^ model seems therefore advisable.

The 10,000 fragments for each superposition were sampled by sampling the fragment starts *s*(*r*) from the distributions in Figure 3 and the fragment lengths *l*(*r*) from a Gaussian with mean 200 bps and standard deviation 80 bps, which is the default setting for fragment length distributions in Cufflinks. These fragments were converted into paired end reads and written to a SAM file [8]. Figures 1 to 4 in the supplement show examples for the different types of coverage obtained by this method.

### 3.2 Experiments with correct annotations

The first set of experiments focused on correct annotations. Thus, the annotations given to the different abundance estimation methods were the same as the ones from which the data were sampled. The Mix^2^ model used in the experiments tied only the mixture weights *β_ij_* and the *p*(*s*(*r*)|*t* = *i, b* = *j*) were Gaussians placed at equidistant positions along the transcript, i.e. *µ_ij_* = *j · l*(*t* = *i*)*/M − l*(*t* = *i*)*/*(2*M)*, where *M* is the number of blocks and *l*(*t* = *i*) is the length of transcript *t* = *i*. The standard deviations of the Gaussians for a transcript were constant, i.e. *σ_ij_* = *l*(*t* = *i*)*/*(2*M)*. Since only two kinds of parameters are estimated, namely the *α_i_* and *β_j_*, this type of Mix^2^ model will be referred to as the 2 parameter Mix^2^ model, and abbreviated as 2p Mix^2^ model. In order to allow for a sensible amount of variability in the fragment bias while avoiding overfitting of the data, 8 blocks were used for every distribution *p*(*r*|*t* = *i*) and the mixtures weights were initialized to be *β_j_* = 1*/*8. The abundances were initialized uniformly, i.e. *α_i_* = 1*/N*, where *N* is the number of transcripts in a gene.

Since the fragment lengths were sampled from the default fragment length distribution used by Cufflinks, the correct fragment length distribution was assumed to be given. Therefore *p*(*l*(*r*)|*t* = *i*) in the Mix^2^ model was a Gaussian with mean 200 bps and standard deviation 80 bps truncated and normalized to the length of transcript *t* = *i*. It would have been possible to estimate *p*(*l*(*r*)|*t* = *i*) from the test data for each gene but the difference between true and estimated fragment length distribution are minor. In general, modifications of *p*(*l*(*r*)|*t* = *i*) in the Mix^2^ model were found to have little influence on the abundance estimates.

The 2p Mix^2^ model was used with and without group tying. For the Mix^2^ model with group tying the transcripts of a gene were placed into one of two different groups separating long from short transcripts. The groups were chosen ad hoc by visual inspection of the gene annotation independently of the fragmentation bias.

Figure 4 shows an example for the convergence of the Mix^2^ model parameters during the course of the EM algorithm, which terminated after 32 iterations. The gene investigated in Figure 4 is the KLK5 gene, whose 5 transcripts are divided into a group containing 4 long transcripts and another group containing a single short transcript. Figure 4(a) shows the convergence of the abundances of the 5 transcripts, where the horizontal lines correspond to the true values. Figure 4(b) and (c), on the other hand, show the convergence of the *β_j_* for the 8 blocks of the group of long transcripts (b) and of the short transcript (c). The blocks are numbered in the 5’ to 3’ direction. Figure 4(b) shows that two blocks at the start of the transcripts and one block at the end of the transcripts have large *β_j_*. This can also be seen from Figure 4(d), where the blue line shows the fragment start distribution *p*(*s*(*r*)|*t* = *i*) that is obtained for the long transcripts after termination of the EM algorithm. In contrast, Figure 4(c) shows that for the group of the short transcript only two blocks at the start of the transcript have large *β_j_*. This is also reflected by the red curve in Figure 4(d), which shows the fragment start distribution obtained for the short transcript at the end of the EM algorithm. Comparing Figure 4(d) and 3(d) shows that the fragment start distributions obtained by the Mix^2^ model closely resemble the distributions from which the data were sampled.

**Figure 4:**
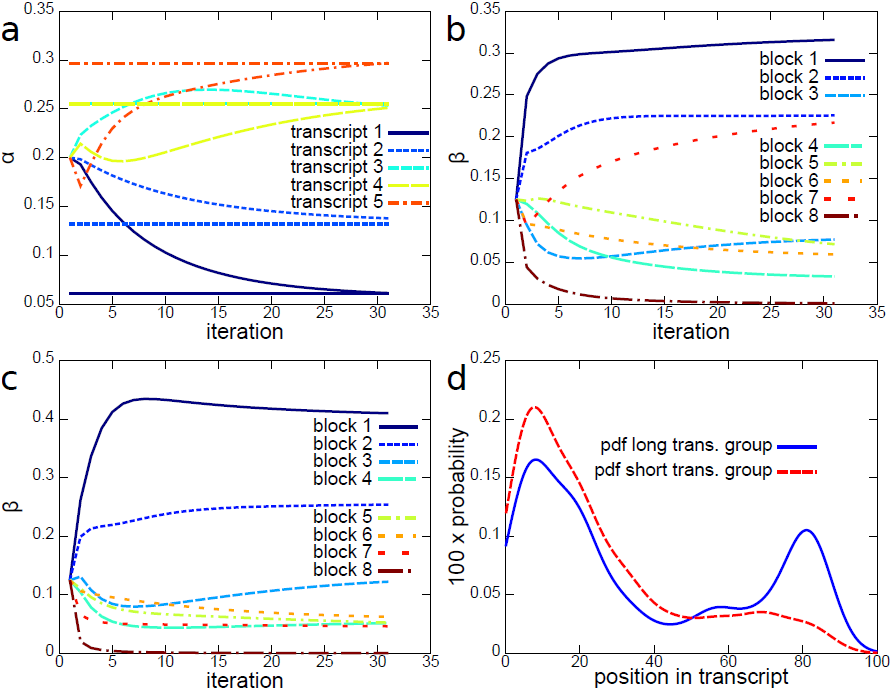
Convergence of the 2p Mix^2^ model parameters in the EM algorithm with correct annotations for the KLK5 gene and fragments with 5’+3’ bias. Figure (a) shows the convergence of the abundances for the 5 transcripts, i.e. the *α_i_*. The horizontal lines in Figure (a) represent the correct abundances. Figure (b) and Figure (c) show the convergence of the *β_j_* for a group containing long transcripts (b) and short transcripts (c). The fragment start distributions in each group are modelled with 8 mixtures, resulting in 8 *β_j_*. Figure (d) shows the probability distributions *p*(*s*(*r*)|*t* = *i*) for the two groups obtained at the end of the EM algorithm. In this example the EM algorithm terminated after 32 iterations.

Figure 5 shows the average L_1_ distance for each gene and each abundance estimation method averaged over the 4 bias models (a) and for each bias averaged over the 7 test genes (b). Thus, as mentioned before, each gene in Figure 5(a) summarizes 800 experiments per abundance estimation method and Figure 5 summarizes 1400 experiments per abundance estimation method. A detailed list of the statistics of the L_1_ distances per gene and per bias model can be found in the supplementary data. In addition, the supplementary data contains boxplots of the differences and L_1_ distances between correct and estimated abundances for the 200 experiments per gene for each combination of bias and abundance estimation.

**Figure 5:**
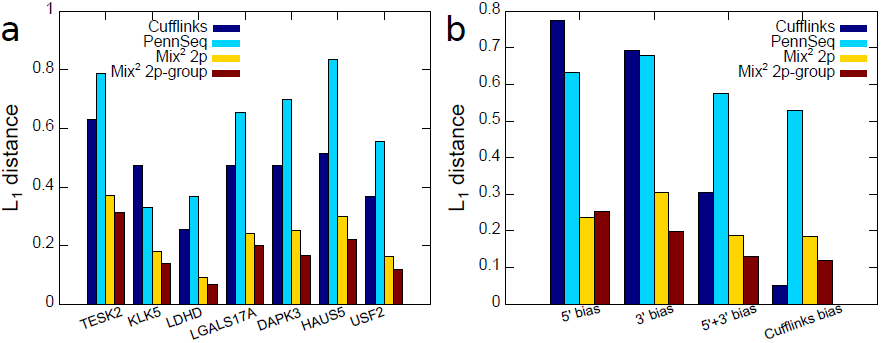
Average L_1_ distances between true and estimated abundances with correct annotations for Cufflinks, PennSeq and the Mix^2^ model with and without group tying. Figure (a) shows the L_1_ distances for each gene in the test set averaged over the different bias types. Figure (b) shows the L_1_ distances for each bias type averaged over the genes in the test set. The numbers visualized in Figure (a) and Figure (b) are given in Tables 13 and 14 in the supplement.

Figure 5(a) shows that averaged over all the 4 biases the 2p Mix^2^ model with group tying yields the smallest error, ranging between 0.07 and 0.31, followed by the 2p Mix^2^ model without group tying. Averaged over all the 4 bias models PennSeq performs worse than Cufflinks on all the genes. Figure 5(b) shows that for the 3’, 5’+3’ and Cufflinks bias the average L_1_ distance for the 2p Mix^2^ model with group tying is smaller than for the 2p Mix^2^ model without group tying. This does not hold for the 5’ bias for which the 2p Mix^2^ model without group tying is more accurate. This is due to the fact that for the 5’ bias in Figure 3(b) the transcript length has little influence on the distribution *p*(*s*(*r*)|*t* = *i*). The introduction of more than one group therefore leads to an unnecessary increase in model parameters which results in overfitting. However, both 2p Mix^2^ models significantly outperform Cufflinks 2.2.0 as well as PennSeq for the 5’, 3’ and 5’+3’ bias models. Not surprisingly, for fragments with Cufflinks bias Cufflinks 2.2.0 outperforms both PennSeq and the Mix^2^ models. In contrast to Figure 5(a), PennSeq seems to give moderate improvements over Cufflinks 2.2.0 for the 5’ and 3’ bias models. For the 5’+3’ and Cufflinks bias, however, PennSeq performs significantly worse than Cufflinks. This seems to suggest that PennSeq performs reasonably well in the presence of strong asymmetric biases, which has also been noted in [5], while its performance deteriorates for biases which are more symmetric and closer to uniform.

### 3.3 Experiments with incorrect annotations

The experiments in this section were conducted with the fragment samples from the last section but with incorrect transcript start and end annotations. For each gene and each of its 200 abundance sets incorrect transcript start and end sites were sampled for all the transcripts of the gene. The sampling was performed by drawing start and end sites uniformly at random from a window surrounding the correct transcript start and end site. This window was the interval ±200 bp around the correct start or end site intersected with the exon on which the correct start or end site was located. For transcript start sites on the first exon of a gene or transcript end sites on the last exon of a gene this window was extended by 200 bp upstream and 200 bp downstream, respectively. The distribution of the differences between correct and incorrect transcript start and end annotation is approximately uniform over the ±200 bp interval but tails off slightly at its start and end due to truncation by short exons. A histogram of the difference between correct and incorrect transcript start and end annotations is shown in Figure 5 in the supplement.

In this section the continuous Mix^2^ model from Section 2.2.2 will be investigated whose mixture components *p*(*r*|*t* = *i, b* = *j*) are given by (9). Only the parameters *α_i_* and *β_j_* as well as the shift and scale parameters, *ν_i_* and *λ_i_*, are estimated and hence the continuous Mix^2^ model in this section will be called the 4 parameter or 4p Mix^2^ model. The extended annotations for the 4p Mix^2^ model, as visualized in Figure 2, are derived by adding 200 bp upstream and 200 bp downstream to the incorrect start and end annotation, respectively. The EM algorithm is executed with 10 different random initializations of the model parameters and from the 10 resulting 4p Mix^2^ models the one giving the highest likelihood to the set of fragments is chosen as the final solution. As in the previous section, 8 blocks are used for *p*(*r|t* = *i*) and the abundances and mixture weights are initialized uniformly. For each group of transcripts in the 4p Mix^2^ model the *µ_j_* are distributed equidistantly, along the longest transcript within the group and the constant *σ_j_* are set accordingly. The initialization of the shift parameter *ν_i_* is varied randomly around 200, which is the relative position of the incorrect transcript start from the start of the extended annotation, and the initialization of the scale parameter *λ_i_* is varied randomly around *l*(*t* = *i*)*/L*, where *L* is the length of the longest transcript within the group.

Figure 6 gives an example for the convergence of the 4p Mix^2^ model during the course of the EM algorithm for the KLK5 gene with incorrect annotations and fragments with 5’+3’ bias. Figures 6(a) and (b) show the convergence of the *λ_i_* and *ν_i_*for the 5 transcripts of the KLK5 gene, whereas Figures 6(c) and (d) show the convergence of the fragment start distributions for transcript 1 and 5, respectively. The dashed and solid vertical lines in Figures 6(c) and (d) represent the incorrect and correct annotations, whereas the dashed curve in these figures represents the initial fragment start distribution before the first iteration of the EM algorithm and the solid curve represents the final estimate after the EM algorithm has terminated. Thus, Figures 6(c) and (d) show that in this example the estimate for the fragment start distributions converges to the correct transcript start and end annotations. In addition, as in the previous section, the shape of the fragment start distributions in Figures 6(c) and (d) resembles the distributions in Figure 3(d) for a long transcript and a short transcript.

**Figure 6:**
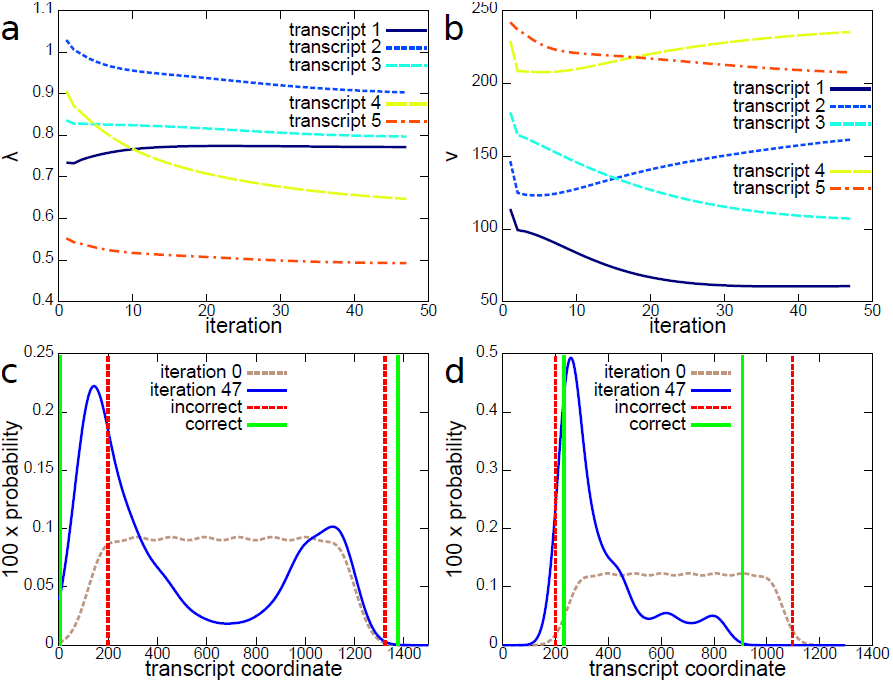
Convergence of the 4p Mix^2^ model parameters in the EM algorithm with incorrect annotations for the KLK5 gene and fragments with 5’+3’ bias. Figures (a) and (b) show the convergence of the *λ_i_* (a) and *ν_i_* (b) for the 5 transcripts of KLK5. Figures (c) and (d) show the convergence of the fragment start probabilities *p*(*s*(*r*)|*t* = *i*) to the correct position for a group containing long (c) and a group containing short transcripts (d). The dashed vertical lines in Figure (c) and (d) are the incorrect transcript start and end annotations, the solid vertical lines are the correct start and end annotations. The dashed curve in Figures (c) and (d) is the initial probability distribution *p*^(0)^(*s*(*r*)|*t* = *i*) with randomly perturbed *λ_i_* and *ν_i_*, whereas the solid curve in Figures (c) and (d) is the final estimate for *p*(*s*(*r*)|*t* = *i*). In this example the EM algorithm terminated after 48 iterations.

For the investigated abundance estimation methods Figure 7 shows the average L_1_ distances between true and estimated abundances, given the incorrect annotations, for each gene averaged over the 4 bias models (a) and for each bias averaged over the 7 test genes (b). Figure 7(a) shows that for each gene the 4 parameter Mix^2^ model with group tying yields the best results, followed by the 2 parameter Mix^2^ model with group tying. As in the experiments with correct annotations shown in Figure 5(a), Cufflinks 2.2.0 outperforms PennSeq with the exception of the KLK5 gene. In comparison to the experiments with correct annotations, however, the difference between Cufflinks and PennSeq is generally smaller. Figure 7(b) shows, again, that for each of the 4 biases the 4p Mix^2^ model performs best, followed by the 2p Mix^2^ model in the case of data with 5’, 3’ or 5’+3’ bias. As expected, for data with Cufflinks bias, Cufflinks 2.2.0 outperforms the 2p Mix^2^ model since the 2p Mix^2^ model can only learn an approximation to the true fragment distribution of Cufflinks 2.2.0. The correction of the inaccurate transcript annotations by the 4p Mix^2^ model, however, is sufficient to yield better results than Cufflinks 2.2.0 even for data with Cufflinks bias. Comparison of the average L_1_ distances for Cufflinks and PennSeq shows that, as before, PennSeq performs slightly better than Cufflinks 2.2.0 on data with 5’ and 3’ bias, while the opposite holds on data with 5’+3’ bias and Cufflinks bias.

**Figure 7:**
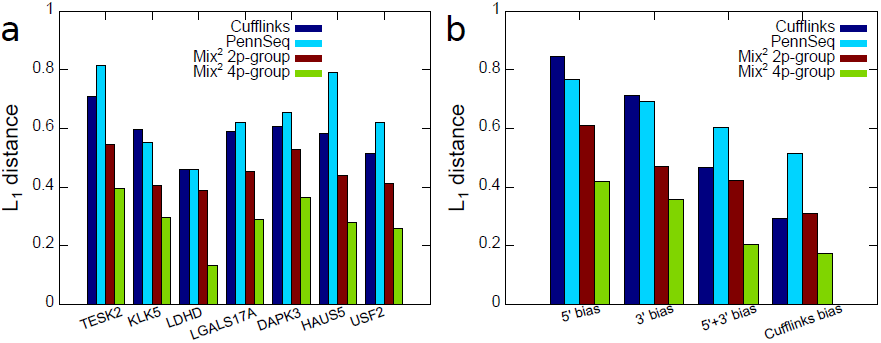
Average L_1_ distances between true and estimated abundances with incorrect annotations for Cufflinks, PennSeq and the 2p and 4p Mix^2^ model with group tying. Figure (a) shows the L_1_ distances for each gene in the test set averaged over the different bias types. Figure (b) shows the L_1_ distances for each bias type averaged over the genes in the test set. The numbers visualized in Figure (a) and Figure (b) are given in Tables 19 and 20 in the supplement.

## 4 Discussion

This article introduced the Mix^2^ model which uses a mixture of probability distributions to model the transcript specific fragment distributions *p*(*r|t* = *i*). Hence, the Mix^2^ model can adapt to the positional fragmentation bias inherent in RNA-Seq data. In addition, the transcript specific shift and scale parameters of the Mix^2^ model enable the correct positioning of the *p*(*r|t* = *i*) within an extended transcript annotation and therefore the correction of inaccurate transcript start and end annotations. Thus, the added flexibility of the Mix^2^ model in comparison to other models yields a more accurate description of RNA-Seq data and therefore more accurate transcript abundance estimates.

The parameters of the Mix^2^ model can be trained efficiently with the EM algorithm and are tied between different transcripts and/or mixture components, depending on the similarity of the transcripts. Tying the parameters guarantees accurate estimates of the transcript specific fragment distributions and therefore accurate estimates of the transcript abundances. This article investigated the tying of the mixture weights alone and in conjunction with the tying of the parameters of the continuous Mix^2^ model.

Experiments were conducted on synthetic data covering 7 genes of different complexity and 4 types of fragment bias. The experiments compared the abundance estimates obtained by Cufflinks 2.2.0, PennSeq and various types of Mix^2^ models. It was shown that for correct annotations the 2p Mix^2^ model with and without group tying outperformed PennSeq on all 4 bias types, while the Mix^2^ models outperformed Cufflinks 2.2.0 on data with 5’, 3’ and 5’+3’ bias. Only on data generated from the Cufflinks model did Cufflinks 2.2.0 yield slightly better results than the Mix^2^ models. For incorrect annotations the 4p Mix^2^ model with group tying outperformed Cufflinks 2.2.0 and PennSeq on all bias types even on data generated from the Cufflinks model.

These results therefore show the superiority of the Mix^2^ model over current state of the art methods as a means for estimating transcript abundances in the presence of positional fragmentation bias and inaccurate transcript annotations.

## Acknowledgement

Andreas Tuerk was partly funded by the Austrian Research Promotion Agency (FFG) under grant number 838191. Gregor Wiktorin was partly funded by the Wiener ArbeitnehmerInnen Förderungsfond (WAFF) under the “Förderung Innovation und Beschäftigung” scheme.

We would like to thank Michael Ante for providing valuable input to the manuscript and testing the Mix^2^ model software. For the latter we would also like to thank Serhat Güler. Special thanks are due to Lexogen’s founder, Alexander Seitz, for encouraging this project.

The methods discussed in this article were filed by A. Tuerk in July 2013 as priority patent application EP13175774.2.

